# Robust and unbiased estimation of the background distribution for automated quantitative imaging

**DOI:** 10.1101/2021.11.09.467975

**Authors:** Mauro Silberberg, Hernán E. Grecco

## Abstract

Quantitative analysis of high-throughput microscopy images requires robust automated algorithms. Background estimation is usually the first step and has an impact on all subsequent analysis, in particular for foreground detection and calculation of ratiometric quantities. Most methods recover only a single background value, such as the median. Those that aim to retrieve a background distribution by dividing the intensity histogram yield a biased estimation in images in non-trivial cases. In this work, we present the first method to recover an unbiased estimation of the background distribution directly from an image and without any additional input. Through a robust statistical test, our method leverages the lack of local spatial correlation in background pixels to select a subset of pixels that accurately represent the background distribution. This method is both fast and simple to implement, as it only uses standard mathematical operations and an averaging filter. Additionally, the only parameter, the size of the averaging filter, does not require fine tuning. The obtained background distribution can be used to test for foreground membership of individual pixels, or to estimate confidence intervals in derived quantities. We expect that the concepts described in this work can help to develop a novel family of robust segmentation methods.

## 1 Introduction

In the age of quantitative high-throughput microscopy, automated image analysis is not only a valuable tool, but a requirement to analyze the ever-increasing amount of collected data. In comparison to manual analysis, not only it provides speed and scalability, but also enables a repeatable objective quantification, which is independent of user bias and can be easily transferable (Ljosa and Carpenter 2009).

An image analysis pipeline deals with transforming the photophysical signal, detected light, into biological observables of interest. While each pipeline is tailor-made for each particular application, they can be usually divided in three major steps: preprocessing, which involves whole image corrections such as illumination correction (Singh et al. 2014), background subtraction, and registration; segmentation and tracking, to identify objects of interest in the image and follow them through space and time; and measuring, to compute observables of interest over these objects, such as object shapes or mean intensities.

In fluorescence microscopy, the objects of interest are labelled with fluorophores, but the collected light not only belongs them, as there are contributions from other sources of signal, such as autofluorescence, out-of-focus fluorescence, stray light or detector noise, collectively referred to as background noise (Waters and Wittmann 2014). Hence, It is important to estimate the background to account for its effect, as it has an impact on all subsequent analysis. Accurate knowledge of the background distribution can improve the segmentation, allowing the detection of dim objects and yielding better defined borders, which in turn impacts the tracking step. More importantly, it can have a major effect in the calculation of some photophysical quantities, as in ratiometric calcium imaging, in which inaccurate background subtraction leads to large errors in the range of 100% (Chen et al. 2006, Shkryl 2020). Additionally, any quantification of its precision would require an estimation of the background dispersion.

Several methods have been developed to deal with the background, which can be broadly classified into intensity thresholding and mathematical morphology methods (Kervrann et al. 2016).

Intensity thresholding methods, such as Otsu’s method (Otsu 1979), assume that objects are brighter than background and calculate a threshold value to separate them. In these methods, background estimation is closely tied with segmentation: the background is the leftover of finding the foreground objects. But they fail when the proportion of foreground and background is not balanced, or in low signal to noise scenarios (Sezgin and Sankur 2004). In the latter case, as foreground and background intensity distributions overlap, no threshold value can fully split them: some background pixels end up classified as foreground and vice versa. In turn, this yields a biased estimation for the background distribution.

As the background is not always uniform, spatially adaptive intensity thresholding methods were developed that apply the same idea locally, over a user-defined window or kernel. Nevertheless, as these methods usually focus on the intensity histogram, they lose all spatial information in the process. In particular, the fact that objects are spatially structured, while the background is usually not.

Other methods are based on mathematical morphology, such as the top-hat filter, h-dome and rolling ball, which is one of the most popular methods to subtract image background. Usually described as rolling a ball under the intensity surface, the rolling ball method consists of applying a grayscale erosion followed by a dilation with a spherical structuring element (Sternberg 1983). But, these methods depend on parameters whose optimal value is closely related to object sizes (Kervrann et al. 2016). Moreover, these methods do not estimate a background distribution, but only a single value at each pixel.

In this work, we demonstrate the first method to recover an unbiased estimation of the background distribution from an intensity image. Through a robust statistical test, it leverages the lack of local spatial correlation in background pixels to select a subset of pixels that accurately represent the background distribution. We show that it outperforms other methods in precision and accuracy. This novel method is both fast and simple to implement, as it only uses standard mathematical operations and an averaging filter. Additionally, its only parameter, the size of the averaging filter, does not require fine tuning.

## 2 Theory

### 2.1 Exploiting the lack of local correlation for background estimation

To generate an estimation of the background intensity distribution, we exploit the lack of spatial structure in the background regions of fluorescence images. In particular, our method transforms an intensity image into values that measure local regularity in the direction of the intensity gradient, which we called the SMO image (fig. 1). Then, assuming a structureless background, it performs a statistical test which selects only a subset of background pixels, but discards almost all of the foreground regions. That subset of pixels form an unbiased estimation of the background distribution. A key aspect of the method is that, being approximately non-parametric, a threshold value for the test can be selected independently of the intensity image, without any a priori knowledge of its background distribution.

**Figure 1.**
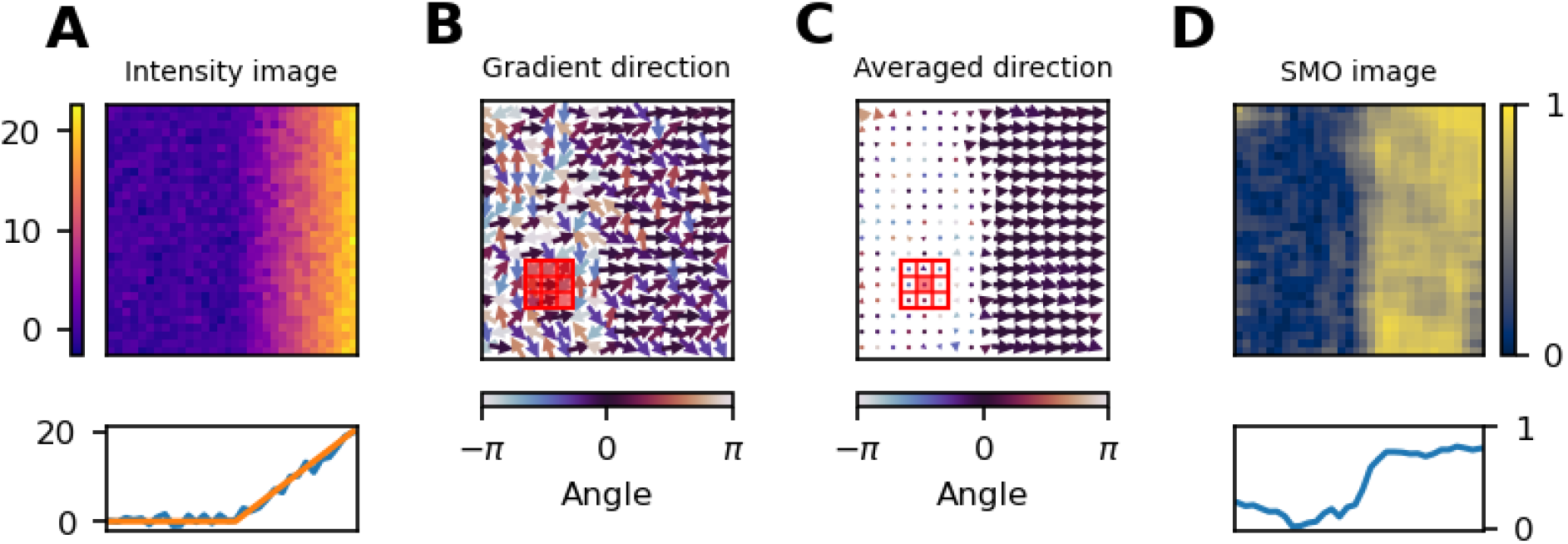
Breakdown of the SIlver Mountain Operator. A) Intensity image simulated as constant intensity followed by a linearly increasing region, plus standard normal noise. On the bottom row, a line profile along a row before (orange) and after (blue) adding noise. B and C) Gradient direction and its local average, respectively, color-coded by angle. The red square indicates the averaging kernel. D) Resulting SMO image, which is the length of the averaged gradient direction.

To demonstrate the fundamentals of the method, we synthetically generated an image consisting of a flat, constant intensity region followed by a linearly increasing intensity region, with added standard normal noise (fig. 1A, see Methods).

For this image, we computed its gradient, a vector measuring of both the rate and direction of greater intensity increase, and then normalized by its length, which results in an image of unit vectors (fig. 1B). Normalizing is what makes this method non-parametric, that is, independent of the underlying distribution. It can be thought of as a multidimensional extension of the Sign test, another non-parametric test, as the normalized gradient in 1D is either +1 or −1, if the intensity increases or decreases from one pixel to the next.

To measure the local variation in gradient direction, we used a local average to calculate a resultant vector for each pixel (red box, fig 1B, and red pixel, fig 1C). The size of the window must be chosen to be smaller than the typical lengths of the gradients.

Finally, each vector was reduced to a scalar value, the SMO value, by calculating its length (fig. 1D). SMO values range from 1, when unit vectors in the averaging windows are aligned, to 0, when they are disordered. As expected from the simulation characteristics, the left and right sides of the SMO image have low and high values, respectively.

In the averaging step, high spatial frequency structures are averaged out and therefore are not detected. For example, a saw-like structure in which the derivative changes direction faster than the averaging kernel will yield a small SMO value and therefore indistinguishable from uncorrelated noise (supp. fig. 1). Therefore, the averaging kernel size must be smaller than the foreground structures, while being large enough to produce a robust estimate (supp. fig. 2).

However, since the normalization step is a nonlinear transformation, the whole process benefits from a previous smoothing step when the gradient is dominated by the high frequency components of the noise, concealing the signal. In these cases, the gradient direction in each pixel will be determined by the noise and therefore will be averaged out in the last step. In turn, this will overestimate the background region (supp. fig. 3).

### 2.2 The SMO distribution is independent of the underlying background intensity distribution

As this method is approximately non-parametric or distribution-free, no a priori knowledge of the underlying background distribution is needed. Additionally, it also implies that SMO values of uncorrelated noise are independent from their intensity values, which is the key for the unbiased estimation of the background distribution.

To demonstrate these properties, we generated uncorrelated noise-only images drawn from different intensity distributions. As no spatial correlation is present, the gradient fluctuates wildly in direction, yielding an SMO image with mostly low values (fig. 2A).

**Figure 2:**
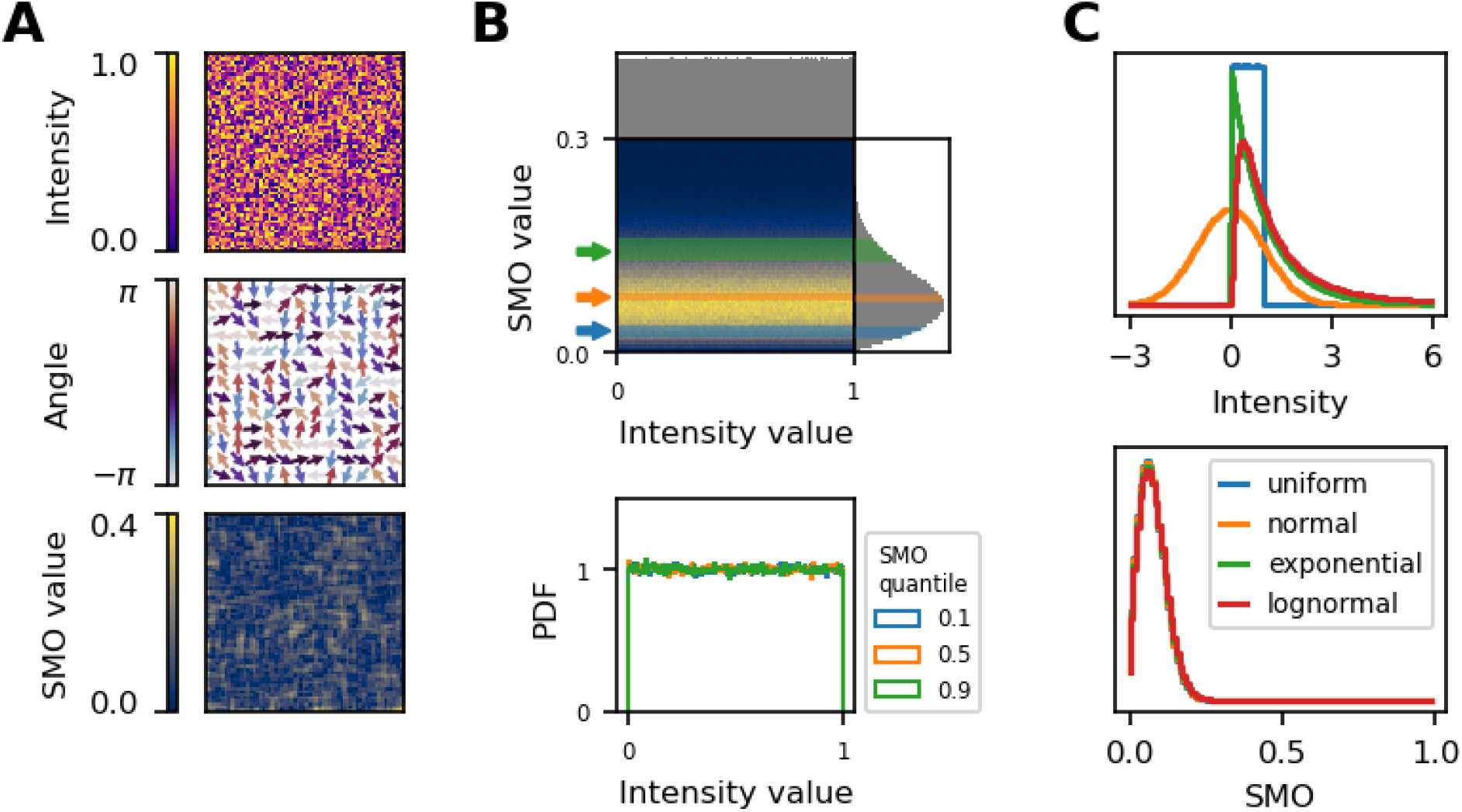
SMO distribution for uncorrelated noise images. A) Simulated random image sampled from a uniform distribution and its corresponding SMO image. B) (top) Joint and marginal distributions of intensity and SMO values. (bottom) Conditional distributions of intensity, corresponding to 0.1-quantile wide slices of SMO ranges as shown above. C) Intensity (top) and SMO (bottom) histograms for uncorrelated noise images generated from different intensity distributions.

The information contained in the SMO values is independent from its corresponding intensity values. Such independence can be seen by the agreement between conditional distributions (fig. 2B, bottom), or by computing its copula from its joint distribution (supp. fig. 5). Each of these conditional distributions corresponds to a 10 percentile-wide slice of SMO values, as depicted in the joint distribution (fig. 2B, top), and yields the same estimation of the intensity distribution. Hence, any subset of SMO values can provide an unbiased estimation for the background intensity distribution.

To showcase that the method is robust against different intensity distributions, we repeated the same procedure for other distributions (fig 2C, top), which produced overlapping SMO distributions (fig 2C, bottom), and independent joint distributions (supp. fig. 5).

In summary, the SMO distribution of a random, structureless image is independent of the underlying intensity distribution, and can be estimated by sampling from any distribution, for instance, a uniform distribution (see Methods, Threshold selection).

### 2.3 Intensity thresholding methods always yield a biased distribution

Due to low photon counts, foreground and background intensity distributions overlap in fluorescence imaging. Hence, intensity thresholding methods are not well suited to the estimation of the background distribution, as it is fundamentally impossible to find a threshold value that splits them. In contrast, our method does not attempt to find the whole background pixels, but only a subset that accurately represents the background distribution.

To simulate this scenario, we generated a circular structure with a bell-shaped intensity profile over a constant background, and added gaussian noise (fig. 3A, see Methods). As the noise amplitude is low compared to the intensity slope, the gradient presents a locally-defined direction in the structure, yielding high SMO values. Instead, low SMO values are assigned to background regions, where the noise level is high compared to the underlying intensity slope (zero), making the gradient direction fluctuate randomly from pixel to pixel.

**Figure 3:**
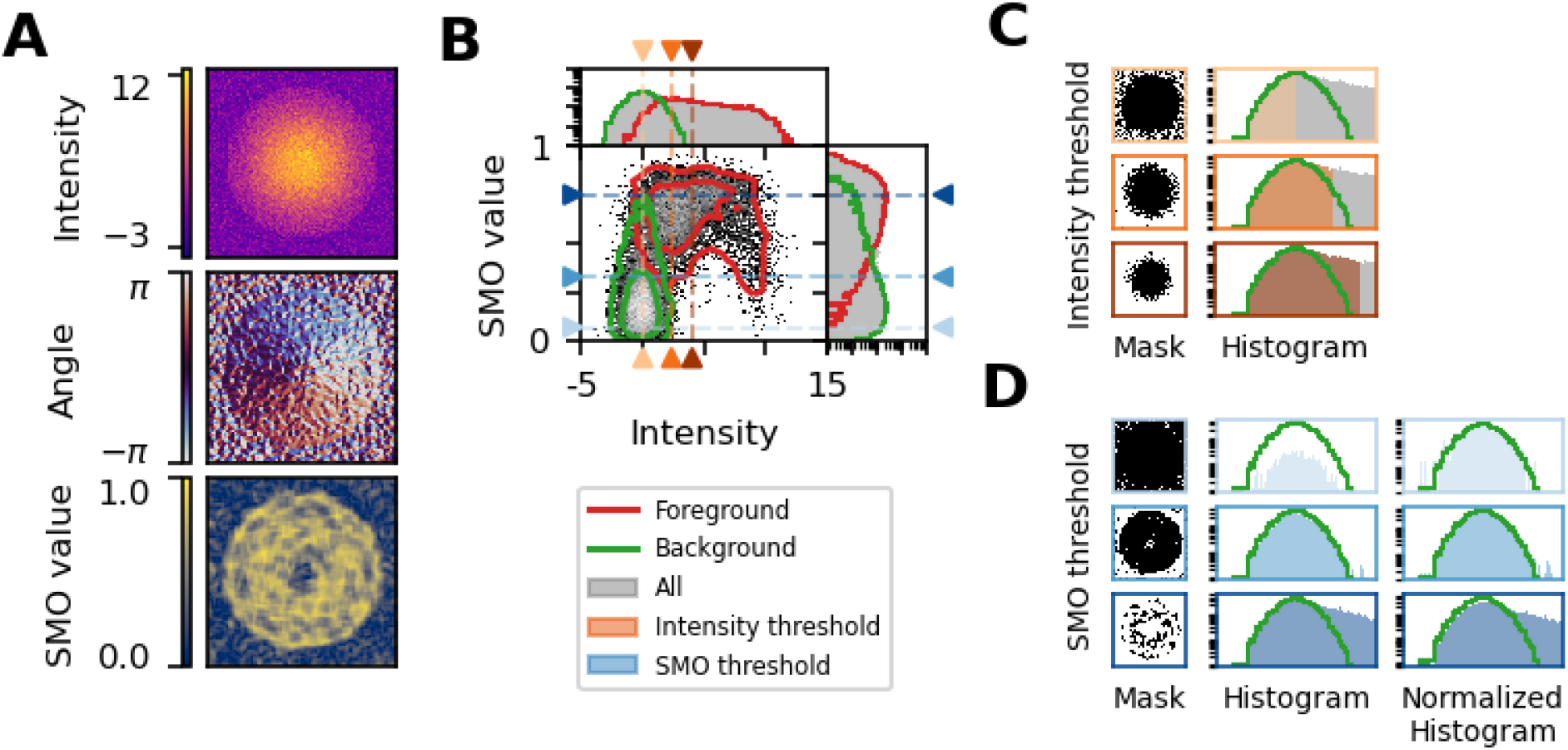
Background estimation in a simulated image. A) Intensity image of a 2D gaussian structure over a constant zero background, plus standard normal noise. Below, the corresponding gradient direction and SMO images. B) Joint and marginal distribution for intensity and SMO, showing separately foreground (red) and background (green) distributions. C and D) Masks and recovered background distributions via intensity (orange) and SMO (blue) thresholding, respectively. Lines and arrows in B show the respective thresholds.

The SMO also assigns low values in the center of the structure, which corresponds to a local maximum in intensity, where the gradient direction changes abruptly. Although the gradient direction has a regularity which distinguishes it from background, as they all point towards the maximum, the average vector is small and yields a low SMO value, which makes it indistinguishable from noise.

Knowing the ground truth, both background and foreground can be visualized separately in joint and marginal distributions (fig. 3B). As they overlap, no single threshold can fully split them, either in intensity or SMO values. Any estimation of the background distribution from an intensity thresholding yields a biased distribution, with either a missing or a larger tail (fig. 3C). Instead, as we have shown that any slice of SMO values samples fairly from a structureless background, we can recover an accurate estimation of the background distribution by using a sufficiently small SMO threshold such as to exclude as much foreground as possible (fig. 3D). As we know a priori the distribution of SMO values for structureless backgrounds (fig. 2C, bottom), we can select an appropriate threshold as a low percentile of that distribution, such as the 10th percentile.

## 3 Results

### 3.1 The SMO matches a manually-recovered background in fluorescence microscopy images

To validate our method in real-world conditions, we used a fluorescence microscopy image of cells transfected with a cytosolic biosensor (fig. 4A, see Methods). It shows a collection of cells with a broad range of intensities spanning the whole dynamic range of the camera. The gradient angle image clearly shows a spatial regularity for each cell, even for very dim ones. In contrast, the background regions lack any spatial structure. Hence, background areas have low SMO values, while cells have mostly high SMO values.

**Figure 4:**
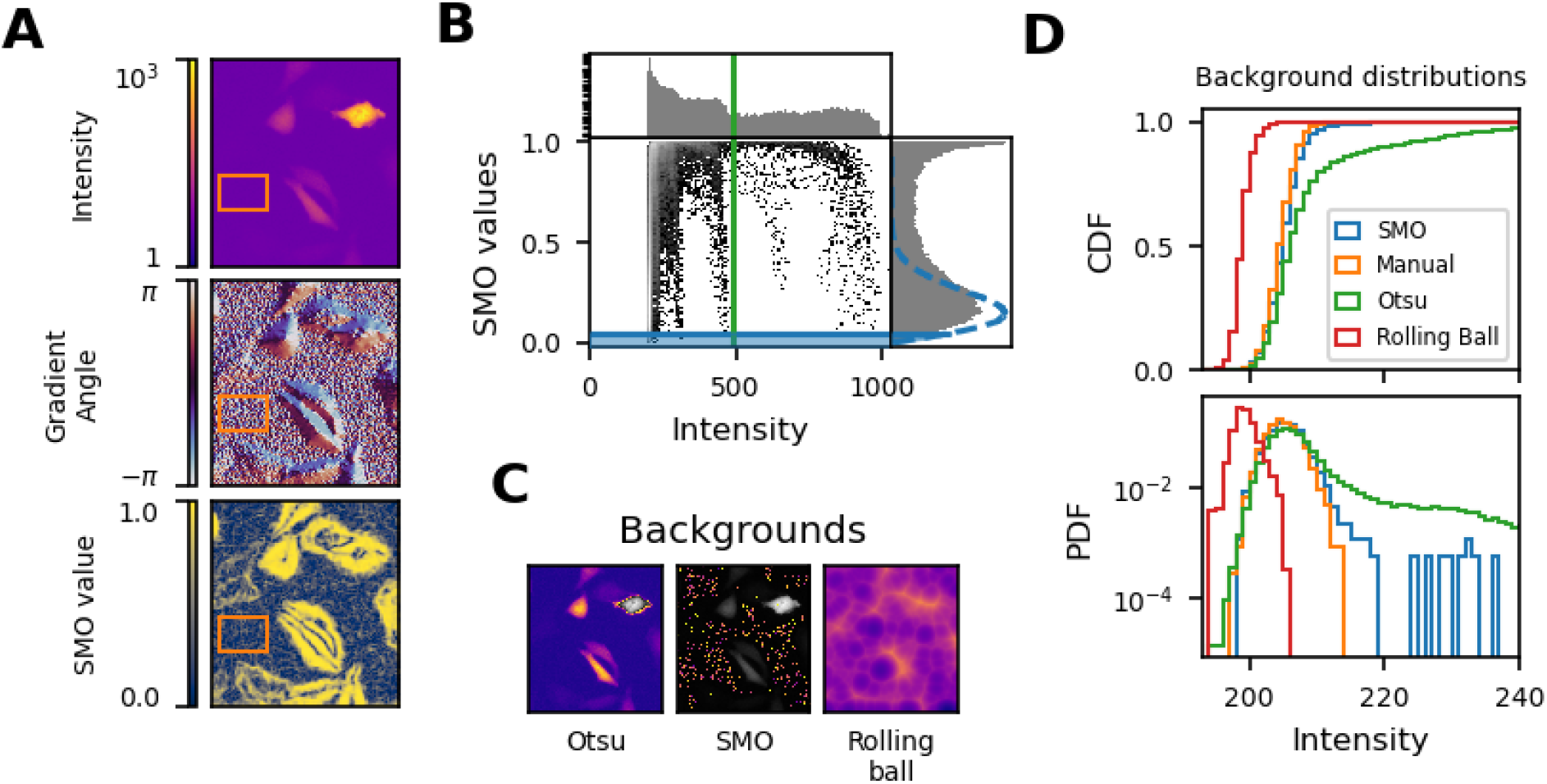
Background estimation in a fluorescence microscopy image. A) Intensity, gradient angle and SMO images. The orange rectangle corresponds to a manually selected background region used as ground truth. B) Grey: joint and marginal distributions of intensity and SMO values. Blue dashed and solid lines: distribution and 5th percentile of SMO values from a simulated structureless image. Green line: Otsu intensity threshold. C) Otsu and SMO and Rolling Ball background estimations. Grayscale regions correspond to excluded areas in the former two methods. D) Comparison of background distributions obtained from ground truth, Otsu, SMO and rolling ball. As cumulative (top) and density (bottom) distribution functions.

As in the simulated example (fig. 3A), cells present low SMO values in their centers, corresponding to local maxima in their intensities. These can also be appreciated as collections of pixels forming an upside-down L shape in the joint distribution of intensity and SMO values (fig. 4B). Likewise, there is no separation between foreground and background in the intensity histogram (fig. 4B, top).

Applying the Otsu intensity thresholding method, it calculates a threshold so high (fig. 4B, green line) as to exclude only the most bright cell (fig. 4C, left). It yields a background distribution which starts to strongly deviate from the manually-selected ground truth (fig. 4A, orange box) at around the 75th percentile (fig. 4D, green and orange curves). As an additional comparison, we included the rolling ball method (Sternberg 1983), which is the standard background correction method in ImageJ. As it is similar to a local minimum filter, it produces a biased distribution in images where the background distribution spans a wide range of intensity values, such as in fluorescence microscopy images. The amount of bias depends on the chosen radius for the ball (supp. fig. 4).

In contrast, thresholding the SMO image at the 5th percentile (fig. 4B, blue line, see Methods for threshold selection) produces a mask that excludes all cells (fig. 4C, middle). While the mask only includes 5% of the background pixels, those constitute an unbiased estimation of the background distribution. In summary, SMO can estimate the background distribution as obtained from manual segmentation in an automatic and robust manner (fig. 4D).

### 3.2 The SMO outperforms standard intensity thresholding methods

Intensity thresholding methods not only yield a biased intensity distribution, but are more sensitive to variations in the foreground intensity distribution. When the intensity histogram is not bimodal, standard intensity thresholding methods fail to produce a good threshold (Kittler and Illingworth 1985). In particular, images from fluorescence microscopy have a strongly unimodal intensity distribution (Baradez et al. 2004).

To evaluate the performance of our method against intensity thresholding, we decided to use fluorescence microscopy images from the Broad Bioimage Benchmark Collection (Ljosa et al. 2012), to achieve a fair comparison without assuming any particular foreground distribution. In particular, we used the BBBC025 dataset, which corresponds to U2OS cells treated with 315 unique shRNA sequences (Singh et al. 2015). From the channel measuring the mitochondria with MitoTracker Deep Red, we selected three images displaying different foreground to background area ratios.

To find a ground truth for the background distribution, we performed a manual segmentation (fig. 5, black curve, see Methods). For images with a low fraction of foreground pixels (fig. 5, left), every method yields the same background distribution up to the 0.8 quantile, since the incidence of foreground pixels wrongly classified as background is not significant.

**Figure 5:**
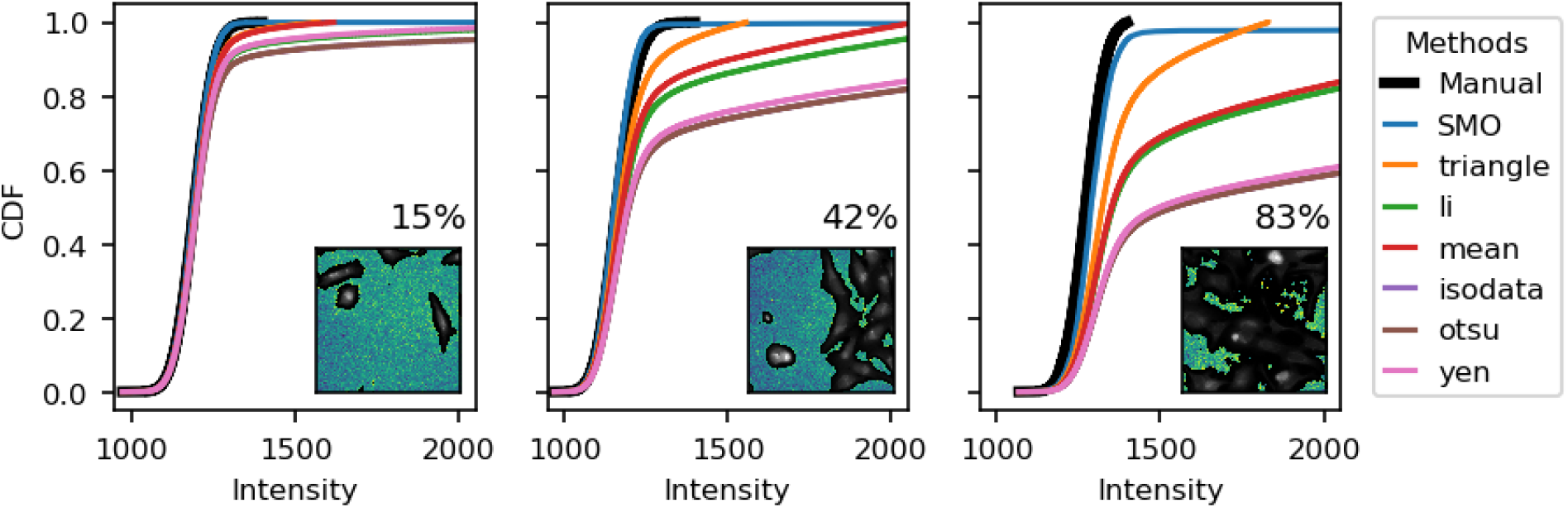
Comparison of background intensity estimation methods for fluorescence microscopy image with differing foreground to background area ratios. The cumulative distribution function (CDF) for different methods is compared against a manually-tuned segmentation (see Methods), shown in the inset. The title of the inset corresponds to the percentage of foreground pixels in the image.

The quality of a blind background estimation method can be assessed by its performance across widely different conditions. As the fraction of foreground pixels increases (fig. 5, center and right), the SMO method (blue curve) remains close to the manually selected ground truth. Instead, in the other methods, up to half of the background distribution actually corresponds to the foreground, leading to a biased median and mean values. Therefore, background correction, which is an essential step in all ratiometric quantifications, will yield a wrong result.

### 3.3 Testing robustness with a high-throughput dataset

To compare the robustness of our method against standard intensity thresholding methods, we used all 3456 fields of view of image set BBBC025, series 37983, from the Broad Bioimage Benchmark Collection (Ljosa et al. 2012). Briefly, eight organelles and cell compartments were labeled: nucleus (Hoechst 33342), endoplasmic reticulum (concanavalin A/AlexaFluor488 conjugate), nucleoli and cytoplasmic RNA (SYTO14 green fluorescent nucleic acid stain), Golgi apparatus and plasma membrane (wheat germ agglutinin/AlexaFluor594 conjugate, WGA), F-actin (phalloidin/AlexaFluor594 conjugate) and mitochondria (MitoTracker Deep Red) (Singh et al. 2015). Image acquisition was performed on 5 different fluorescent channels, as described in Gustafsdottir et al. 2013, which we identify according to their filename.

As all images in the same channel and acquisition settings have the same underlying background, we used the variation in median background intensity for each channel as a proxy for robustness. For each image in the dataset, we computed its background distribution and its median (fig. 6A, blue histogram and orange line). For each method, we built the distribution of median backgrounds (fig. 6A, orange histogram), from where we interpret a smaller variation in median backgrounds as a more robust method.

**Figure 6:**
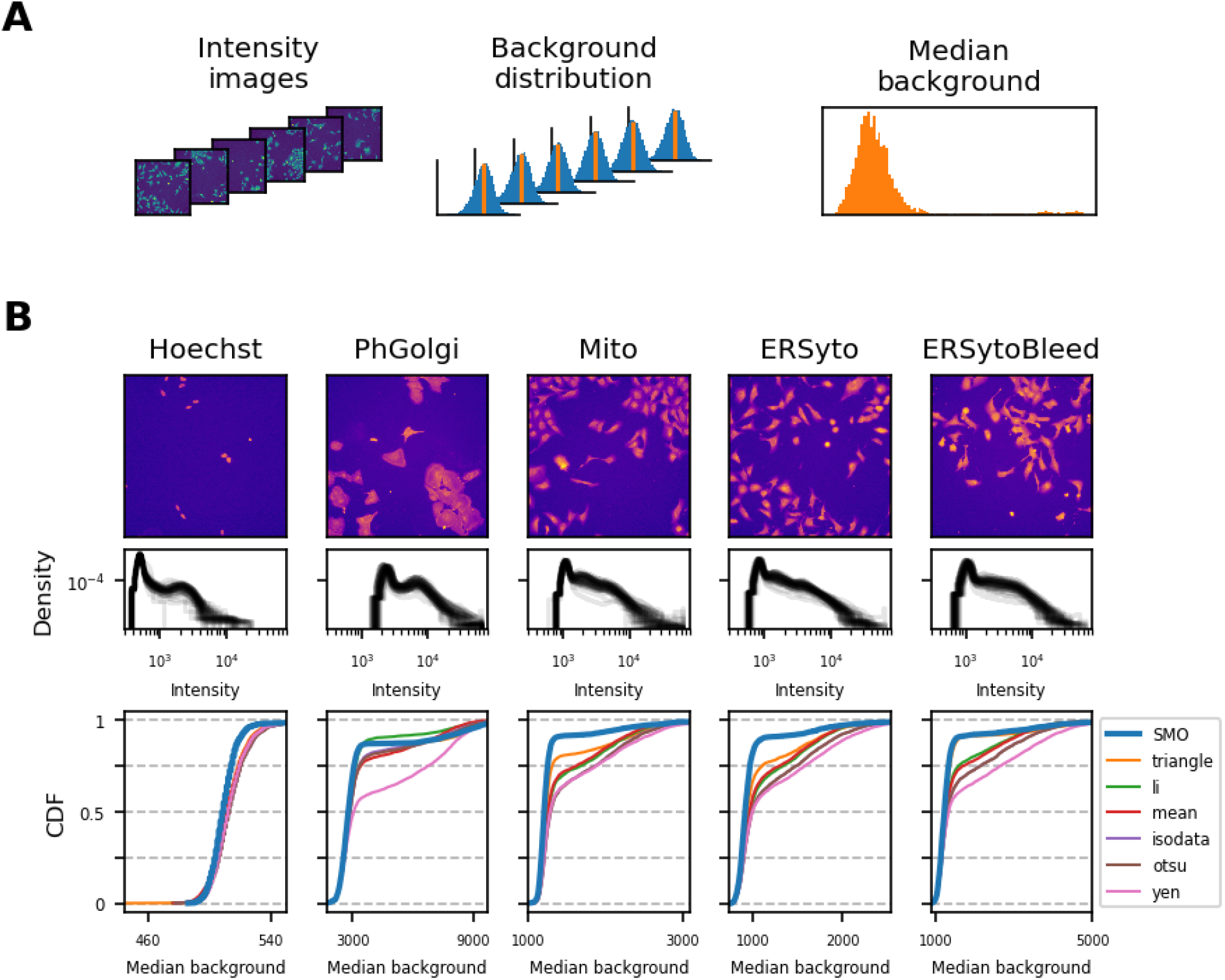
Comparison between methods for estimation of the median background for the dataset 25 of the Broad Bioimage Benchmark Collection. A) For each image, we estimated the background distribution (blue), computed its median (orange). Then, we built the distribution of median backgrounds for all images. B) On the top row, a typical image is shown for each channel. On the next row, a sample of image histograms for some of the images. The third row shows the cumulative distribution function (CDF) for the median backgrounds of all the dataset.

As expected, in the Hoechst channel (fig. 6B, first column), which marks the DNA, all methods behave properly, presenting a low variation, as indicated by the steep cumulative distribution function (CDF, fig 6B, bottom row). As the fluorescence is localized in the nucleus and has a low cell to cell variation, it gives rise to a bimodal distribution (fig. 6B, middle row), for which the intensity thresholding methods perform well. While the variation is small, there is a bias between the intensity thresholding methods and the SMO, as depicted by the shift in the blue curve. A similar behaviour is observed for the PhGolgi channel, which also shows a bimodal distribution. But in this case, Yen’s method shows a marked deviation in the median background for almost half of the dataset, as illustrated by its change of trend around the 0.5 quantile (fig. 6B, pink curve).

For the case of the Mito, ERSyto and ERSytoBleed channels, which do not exhibit a marked bimodal distribution, the variation in median background is much more pronounced for the intensity thresholding methods. All methods deviate around the 0.5 and 0.8 quantiles, yielding higher estimates for the median background. A subsequent analysis shows that this deviation is highly correlated with the foreground to background area in the images (supp. fig 7).

## 4. Discussion

### 4.1 Overview

We developed a statistical test that exploits the lack of correlation between neighboring pixels in the background to select a subset of pixels that accurately represent the background intensity distribution. Furthermore, being a non-parametric test, no a priori knowledge of the background distribution is needed.

We applied this method to a low signal to noise synthetic image, where we showed that intensity thresholding yields a biased distribution, while a range of SMO thresholds provide an unbiased estimation of the background distribution. Also, we applied it to a fluorescence image of a cytosolic biosensor, where it obtained a more accurate background distribution than the rolling ball method. We also found that SMO is more robust than any other method as evidenced by measuring the median of the background in a large and diverse dataset. While some large datasets include a segmentation ground truth, we have found a large false positive rate in the identification of background pixels (e.g. dim cells) and therefore it is not reliable to test for robustness.

In contrast to methods that only estimate a single background value, our method obtains an accurate estimate of the background distribution. With this knowledge, it is possible to evaluate the bias and uncertainty in the calculation of biological observables, in particular in the case of ratiometric calculations (Kalinin et al. 2008).

Our method requires that structures present a gradient, i.e. that objects do not have flat intensity profiles. This is a common feature in fluorescence images of most cell types, particularly when there is not a clear separation between foreground and background, and therefore other methods that rely on finding borders fail. It relies on calculating a measure of the local average in gradient direction and has only one free parameter: the size of the averaging kernel. As long as it is smaller than the typical foreground structure, It doesn’t require fine-tuning, being robust across a wide range of values. Therefore, it’s only necessary to tune it for the first image.

### 4.2 Perspective

A possible improvement to the SMO could replace the local average of gradient directions with a filter that distinguishes local maxima of intensity from uncorrelated background. For instance, specialized tests for circular uniformity (Pycke 2010) might improve discrimination of intensity valleys, albeit with a slower run time. A simpler way is to combine SMO with other established methods. For example, the morphological operations could be used to exclude both pixels from the ridge of cells and areas near their borders, which would improve the background estimation, removing the spurious tail to the right (supp. fig. 6).

The method assumes that the background distribution is the same for each pixel, that is, independent of the position. In cases in which assumption is not fulfilled, for instance, a non-homogeneous illumination, the method can be applied locally, as long as the slope of the background is smaller than changes due to noise. For example, a smooth surface could be fitted to the intensity values selected with the mask obtained via SMO thresholding.

Finally, the SMO image could be used as an input to segmentation algorithms, such as those relying on global thresholds, or machine learning based ones that use several input filters. Furthermore, using the estimation of the background distribution, the intensity image can be transformed to an image of probability of belonging or not to background.

## 5 Methods

### Software

all image and figure generation was made with the Python scientific stack (NumPy, SciPy, matplotlib, scikit-image). Code to reproduce all figures and a Python implementation of the Silver Mountain Operator is available at https://github.com/maurosilber/SMO. We also provide plugins for napari, CellProfiler and ImageJ/FIJI.

### Silver Mountain Operator (SMO) calculation

starting from an intensity image I,

1. (optionally) apply a smoothing filter
2. calculate the image gradient, *∇I*
3. normalize each gradient vector with its length, *∇I*/ ∥ **∇I** ∥
4. apply a moving average to the normalized gradient
5. compute the length of each resulting average vector

In particular, the implemented SMO uses a gaussian filter from SciPy’s ndimage module, calculates the gradient with NumPy’s function gradient, and vector lengths with the euclidean norm.

### Threshold selection for SMO

generate an image sampled from a uniform distribution, calculate its SMO image, and compute a desired percentile. For instance, the 5th percentile to include only 5% of the background. The SMO calculation for the randomly sampled image has to use the same smoothing and averaging filter as the ones used for the target intensity image. Also, it is important to exclude saturated intensity pixels from the resulting SMO mask.

### Simulated structure

a 2D gaussian profile with a peak amplitude of 10 and 100 pixels of radius over a zero background, plus standard normal noise. The total image size was 200×200 pixels.

### Sample preparation and image acquisition

samples correspond to HeLa cells transfected with CASPAM, a cytosolic biosensor described in Habif et al. 2021. Imaging was performed by means of a custom built setup as described in Squire et al. 2004 and Vilar et al. 2009. Briefly, it was composed of an Olympus IX81 inverted microscope (Olympus, Germany) equipped with a MT20 illumination system, an Orca CCD camera (Hamamatsu Photonics, Japan), and a 20X 0.7 NA air objective. The CellR software (Olympus, Germany) was used to acquire images.

### Image analysis for BBBC025

for each of the intensity thresholding methods, the background for each image was estimated as all pixels below the computed threshold. For the SMO method, an averaging window of 7 pixels was chosen, with no previous smoothing. The threshold was chosen as the 5th percentile as described above (Threshold selection for SMO). As these methods assume a homogeneous background, the images were cropped from 1080×1080 pixels to the middle 432×432 pixels region.

## 6 Funding

This work was supported by the following grants: PICT 2014-3658; PICT 2013-1301; Max Planck Gesellschaft Partner Group.

## Notes

### Competing Interest Statement

The authors have declared no competing interest.

