## Supplementary Information for "Robust and unbiased estimation of the background distribution for automated quantitative imaging"

### High-frequency saw-like structure yields small SMO values

As the derivative changes direction faster than the averaging kernel, it yields a null SMO value and therefore indistinguishable from uncorrelated noise.

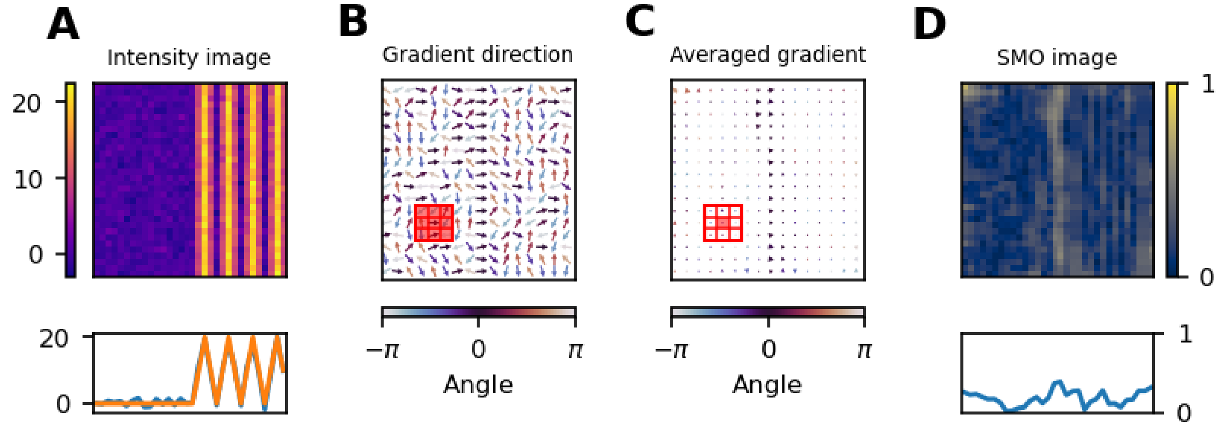


**Supplementary Figure 1. Breakdown of the SIlver Mountain Operator. A) Intensity image simulated as constant intensity followed by a saw-like region, plus standard normal noise. On the bottom row, a line profile along a row before (orange) and after (blue) adding noise. B and C) Gradient direction and its local average, respectively, color-coded by angle. The red square indicates the averaging kernel. D) Resulting SMO image, which is the length of the average gradient.**

### Robustness of for different averaging kernel sizes

The choice of averaging kernel size does not have a great impact on the recovered background distribution, as long as it is smaller than the foreground structures, since they would be averaged out. Nevertheless, larger kernels do produce a slightly better estimate, as more of the neighbourhood is considered to decide if a pixel is part of the background or not.

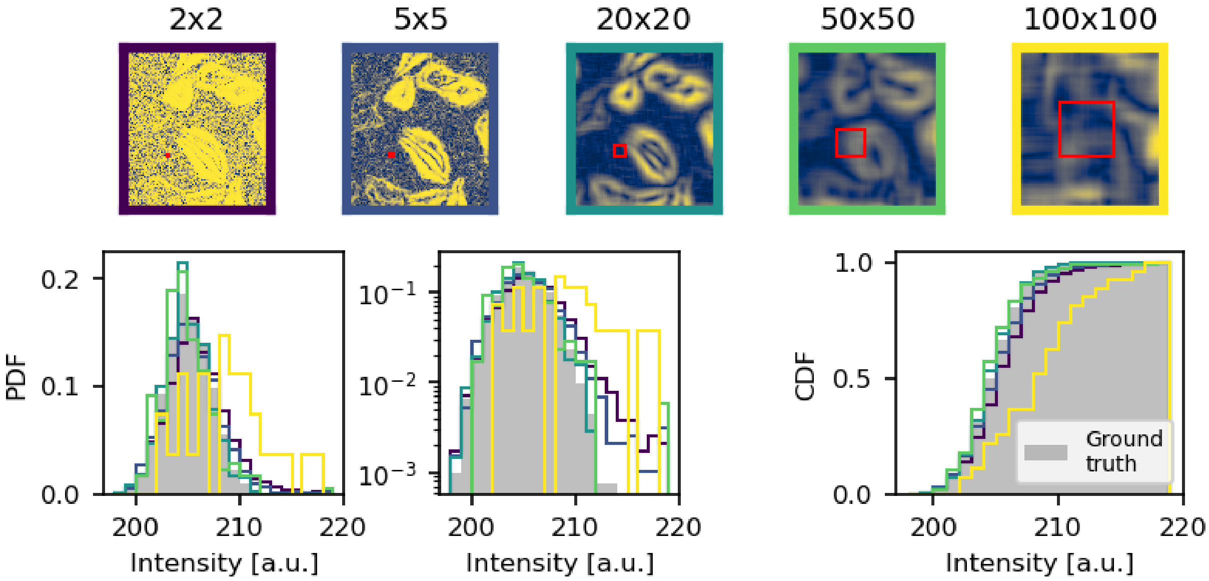


**Supplementary Figure 2. The SMO is robust to changes in averaging kernel size. Top row: SMO images for different kernel sizes, which are shown to scale as red squares. Bottom row: background distributions estimated for each kernel size, compared to a manually selected region as ground truth.**

**A previous smoothing filter improves the performance of SMO**

The SMO benefits from applying a previous smoothing filter to the intensity image, as it can better distinguish the intensity gradient from noise. Note that the thresholded mask obtained from the SMO image is applied to the original intensity image, and not the smoothed one.

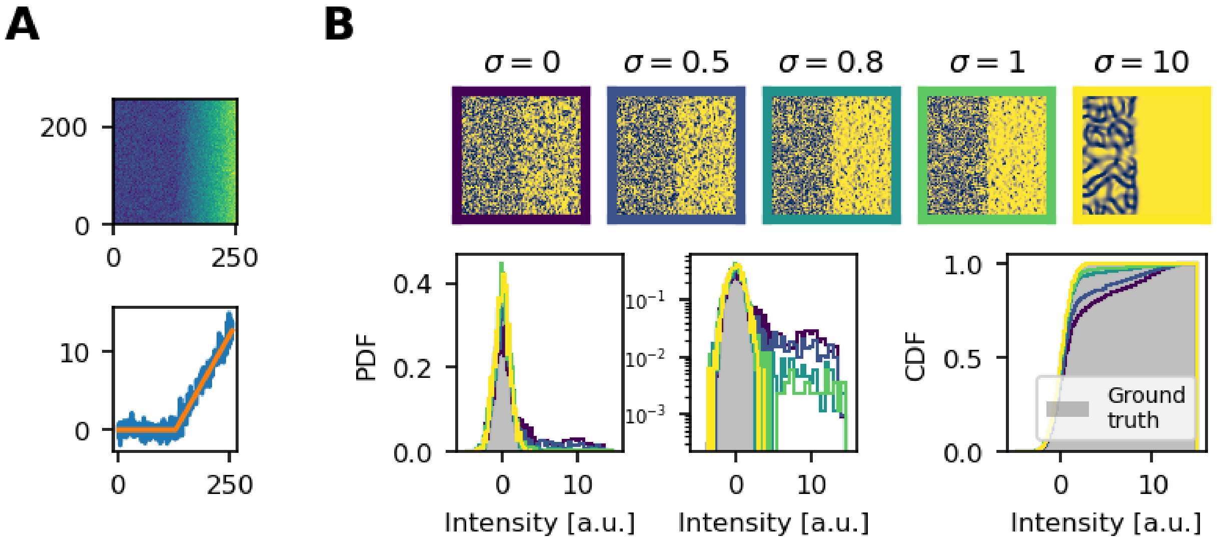


**Supplementary Figure 3. (A) Intensity image simulated as a constant region (background) followed by a linearly increasing region (foreground), plus standard normal noise. (B) Top row: SMO images after applying a gaussian filter of size σ. Bottom row: background distributions estimated, compared to the ground truth (left side of the image).**

**The Rolling Ball method recovers a biased background distribution**

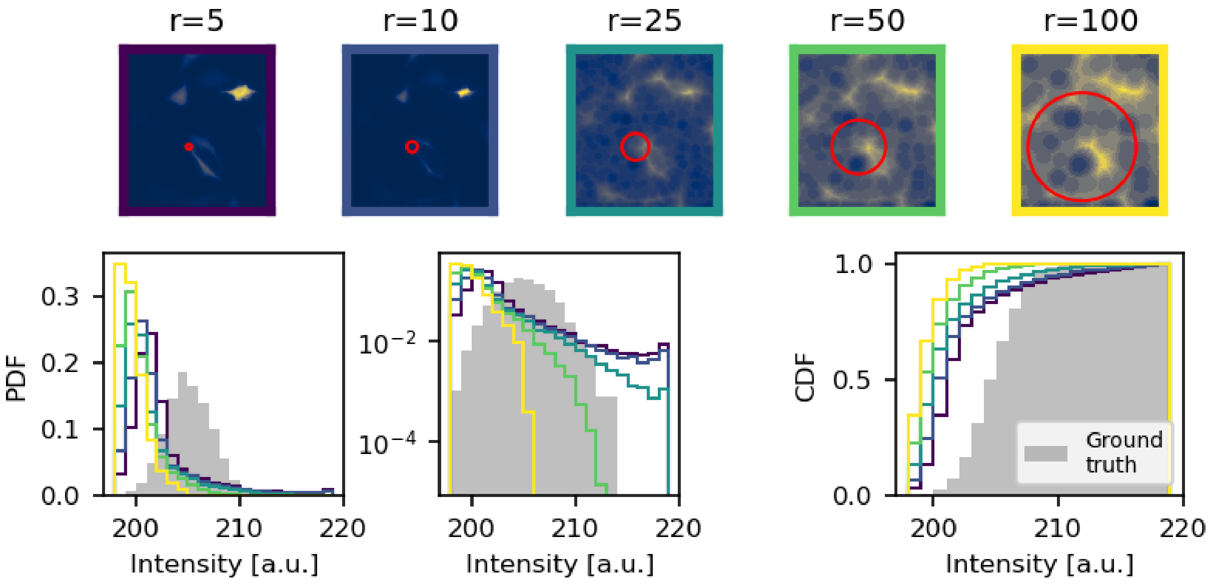


**Supplementary Figure 4. The rolling ball method applied to the intensity image of figure 5. Top row: background estimation different radii $r$. Bottom row: background distributions estimated, compared to a manually selected region as ground truth.**

**SMO values are independent from intensity values for all distributions**

A uniform copula shows independence between intensity and SMO values for the different noise distributions.

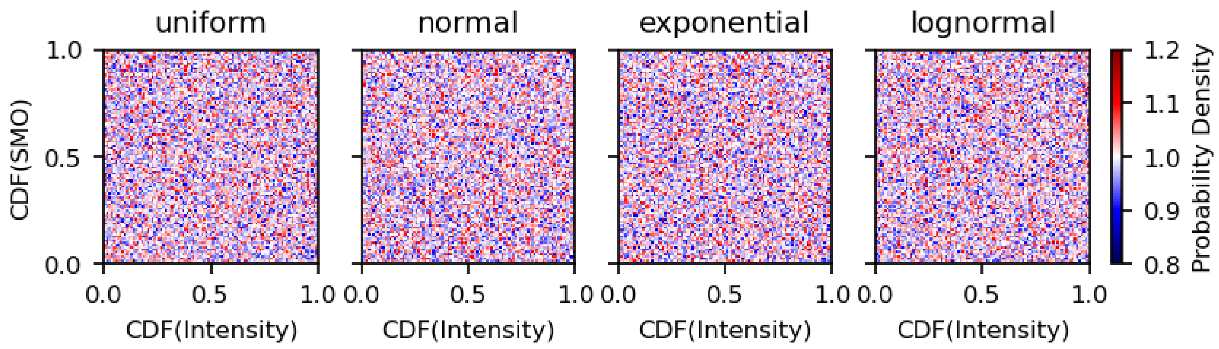


**Supplementary Figure 5. Copulas of the joint distribution of intensity and SMO values for different noise distributions.**

**Refinement of the SMO mask with morphological operations**

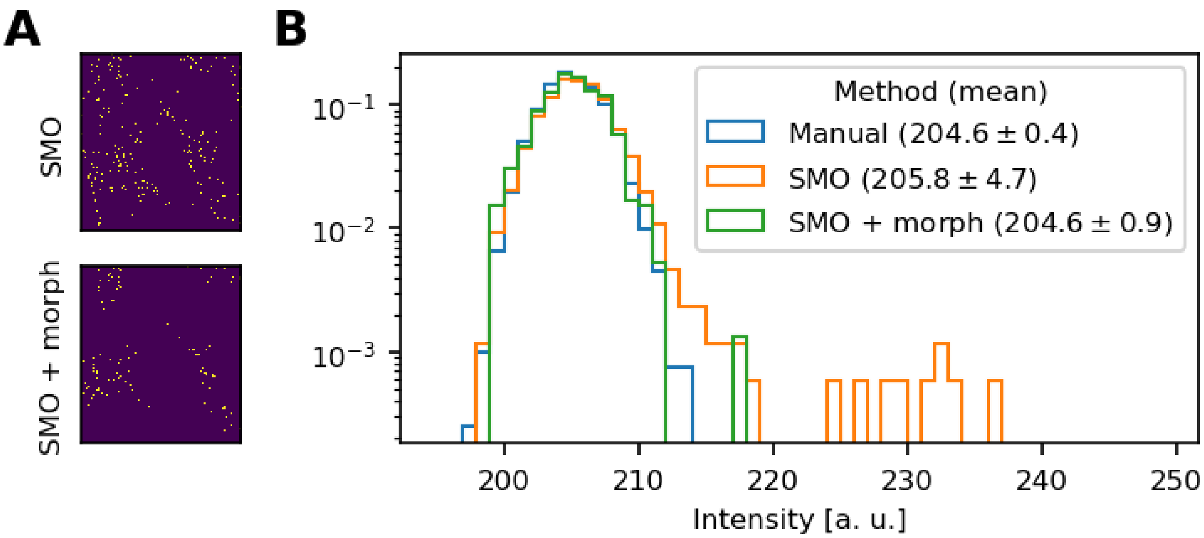


**Supplementary Figure 6. Refining the SMO mask with morphological operations improves the background estimation. (A) Top: Original SMO mask. Bottom: refined with morphological operations to exclude points near to cell borders. (B) Background distributions estimated, compared to a manually selected region as ground truth.**

### Correlation between background to foreground area ratio and method failure in BBBC025

When the number of background pixels is small compared to the foreground ones, all methods fail, estimating a higher value for the median background. While there is not much difference in the Hoechst channel, in the Mito, ERSyto and ERSytoBleed, all methods show a lower breaking point than the SMO method.

As we had no segmentation available as a ground truth, we used a manually selected threshold per channel to segment into foreground and background and calculate area ratio. Hence, it is not the true ratio, but a proxy of it. The chosen thresholds were:

| Hoechst | PhGolgi | Mito | ERSyto | ERSytoBleed |
|---|---|---|---|---|
| 540 | 3500 | 1300 | 1100 | 2000 |

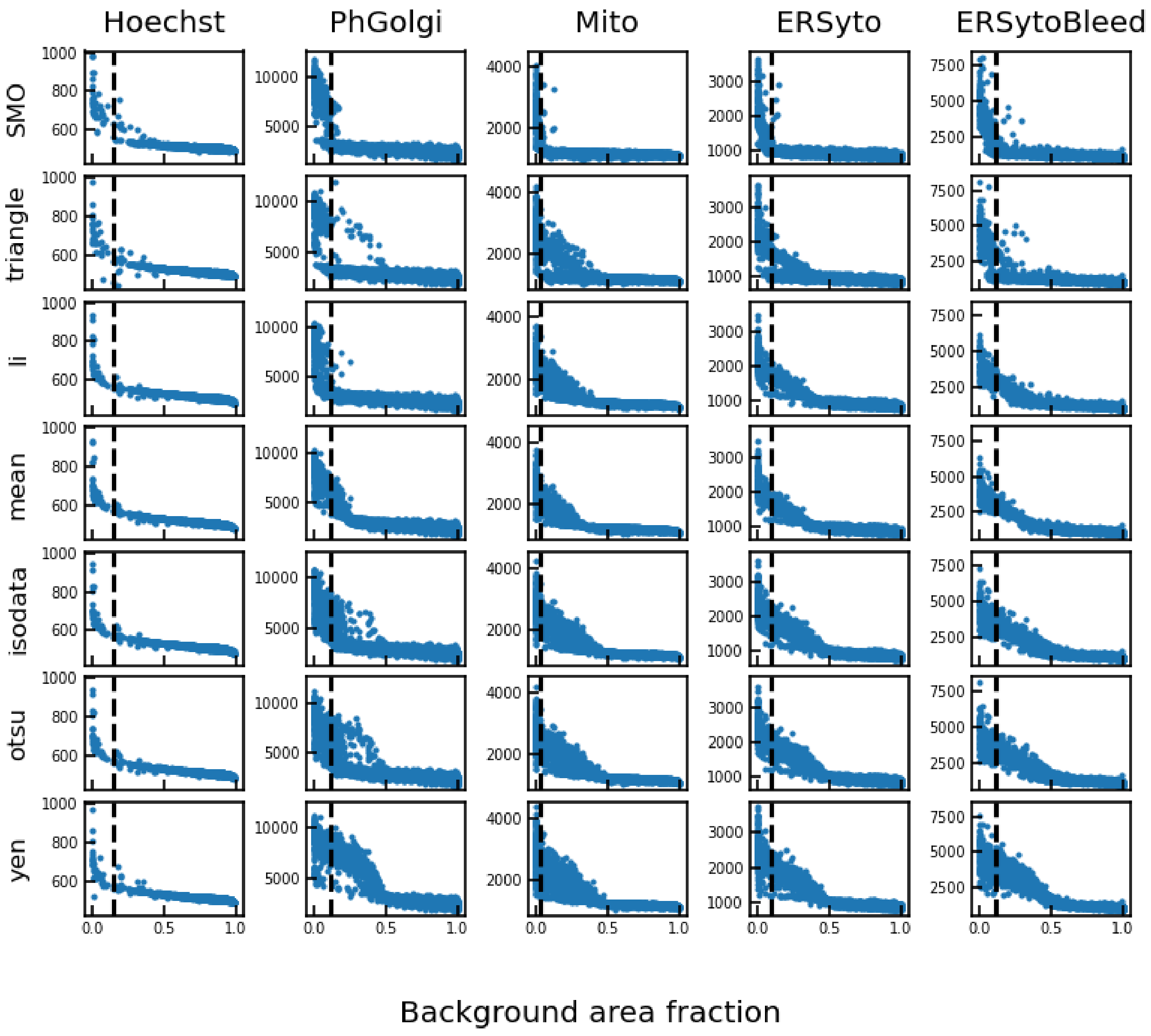

**Supplementary Figure 7: Correlation between foreground-to-background area ratio and bias in median background intensity. The vertical dashed line marks the breaking point for the SMO method.**

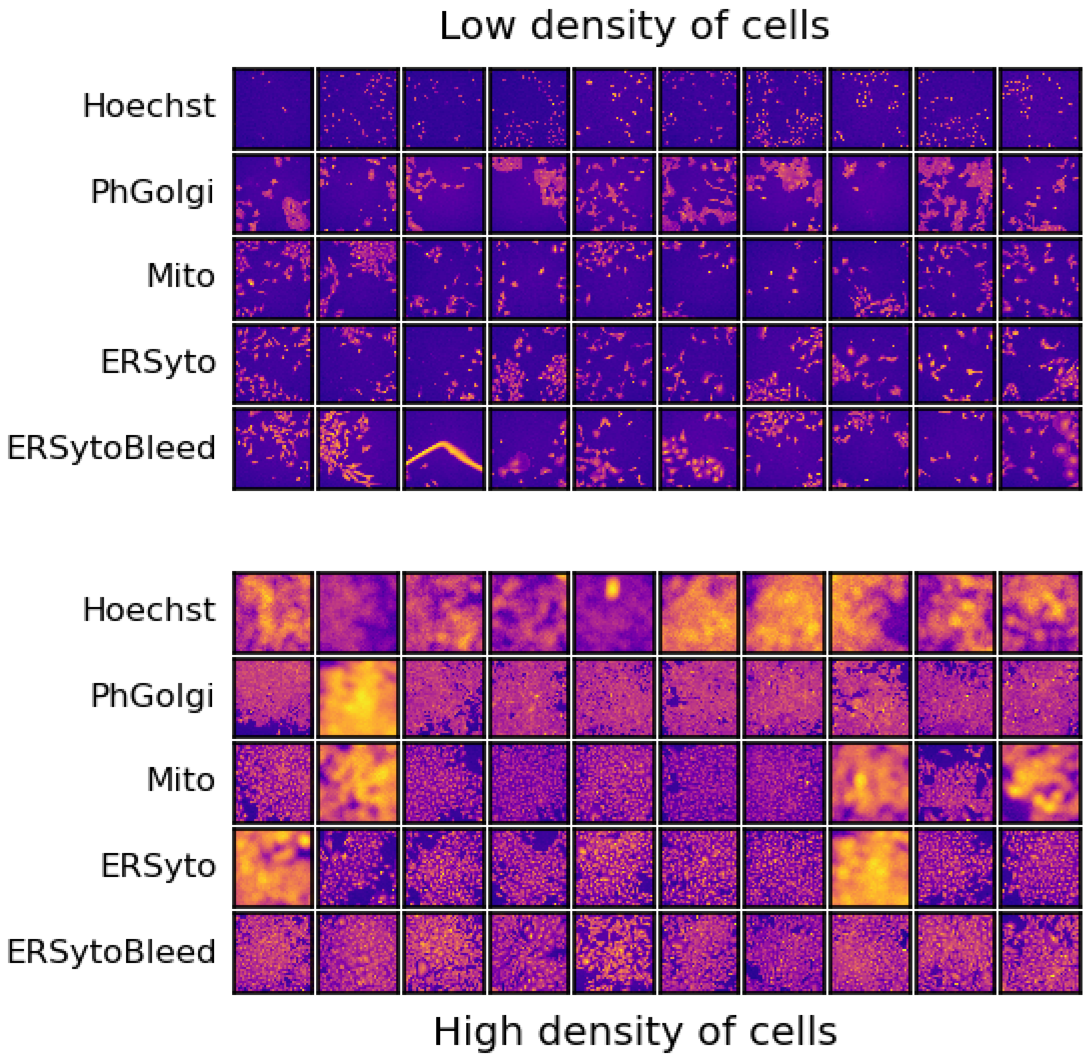


**Supplementary Figure 8: Sample images of low and high density of cells. Top: sample images selected such that all methods reported a median value between the 10 and 20 percentile of the median background distribution. Bottom: all methods reported median background greater than the manually-selected thresholds.**
